# Dataset of the construction and characterization of stable biological nanoparticles

**DOI:** 10.1101/2020.08.23.263590

**Authors:** Romina A. Gisonno, M. Alejandra Tricerri, Marina C. Gonzalez, Horacio A. Garda, Nahuel A. Ramella, Ivo Díaz Ludovico

**Affiliations:** Instituto de Investigaciones Bioquímicas de La Plata (INIBIOLP), Argentina; Facultad de Ciencias Médicas, Universidad Nacional de La Plata. Calle 60 y 120, La Plata. CP 1900, Argentina

**Keywords:** Apolipoprotein A-I, BS^3^ Crosslinker, Lipid-Binding, gradient gel electrophoresis, nanoparticles

## Abstract

We suggest that the structural flexibility is key for certain proteins in order to fulfill functions that are required to interact with biological membranes, and that intra-chain chemical crosslinking may result in a different arrangement of protein with lipids. As interaction with biological membranes and lipids is a function attributed to many proteins in circulation, we intended to characterize an experimental design that helps in the study of many biological protein structures and their function. But in addition, by introducing intra-chain crosslinking, we obtained discoidal nano platforms that are stable under different conditions of temperate and time incubation. These platforms might be an excellent model to employ as biological carriers of intrinsic or external molecules. Thus, data shown here clearly strengthen the usefulness of an easy, accessible and inexpensive tool not only to study protein-lipid interactions, but to be used in different biological fields that require the transport of organic compounds.

## Specifications Table

|  |  |
| --- | --- |
| **Subject** | Biochemistry and Biophysics |
| **Specific subject area** | Biophysics (lipid/protein assay)<br>Protein structure analysis protein lipid Assays |
| **Type of data** | Image<br>Graph<br>Figure |
| **How data were acquired** | The lipid-protein interaction study was carried out through native gels. All these were run in a Mini-PROTEAN® (BIO-RAD Tetra Vertical Electrophoresis Cell for Mini Precast Gels, 4-gel #1658004). Gels were quantified in ImageJ software version 1.51 j8.<br>Micellization assays were monitored alternatively in a Helios Beta: Single beam, quartz coated (Thermo) and in a UV-visible Agilent Cary 8454. (Agilent Technologies), spectrophotometers. Curves were obtained using SigmaPlot 12.0 (Systat Software, Inc. SigmaPlot for Windows). Statistical assays were performed on GraphPad Prism version 8.0.0 for Windows, (GraphPad Software, San Diego, California USA, www.graphpad.com).<br>The relative size of the particles was estimated by both electrophoresis and size exclusion chromatography (SEC) using a Merck-Hitachi L6200 Intelligent pump. Samples were eluted through a 6 HR column (Pharmacia) and an UV-VIS detector (Merck-Hitachi L4200). Particle morphology was characterized by transmission electron microscopy (TEM) on a JEOL-1200 EX. |
| **Data format** | Raw<br>Analyzed<br>Filtered |
| **Parameters for data collection** | Data was collected by three or five independent experiments using recombinant proteins and commercial lipids. |
| **Description of data collection** | Each DMPC curve-point was normalized to absorbance at minute 0 at $\lambda_{325\text{nm}}$.<br>Quantification of lipoproteins was performed by scanning the lane density after imaging the entire gels, by using ImageJ software version 1.51 j8. |
| **Data source location** | Institution: Instituto de Investigaciones Bioquímicas de La Plata (INIBIOLP)<br>City: La Plata, Buenos Aires<br>Country: Argentina |
| **Data accessibility** | With the article<br>Direct URL to data: Díaz Ludovico, Ivo; Ramella, Nahuel A. (2020), "Dataset of the construction and characterization of stable biological nanoparticles", Mendeley Data, V2, doi: 10.17632/rgk9n9wt3d.2 |

## Value of the Data

- Data show a robust experimental approach to construct stable nanoparticles which may act as platforms to be used as carriers of drugs or biological compounds.
- These data may benefit the extended field of either basic or applied biochemistry research, as it may give information on the protein structure-function relationship. Its simplicity and low cost allow being extensively used.
- Nanoparticles as vehicles for drugs are a potent tool for the pharmaceutical and cosmetic industry, as well as for basic research. This design may be combined with Mass Spectrometry, Flow cytometry or fluorescence techniques to study the efficiency of their interactions with biological membranes.
- Data show a well-defined methodological design to evaluate the interaction of flexible proteins with different lipid microenvironments and the importance of a structural constraint as induced here by intra-chain crosslinking.

## Data Description

The efficiency of protein solubilization as lipid complexes and the effect of crosslinking in this behaviorwas evaluated initially by using detergent-mediated interaction. In this regard, reconstitution of apoA-I into discoidal complexes may be attained by using sodium cholate, an amphipathic molecule that, due to its flat-shaped structure was shown to promote the formation of bilayer-like complexes [1]. In this regard, apoA-I with the native sequence (Wt), either unmodified or crosslinked (Wt+BS^3^) was incubated with dimyristoyl phosphatidylcholine (DMPC) multilamellar vesicles (MLV) at a 40:60:1 lipid/sodium cholate/protein molar ratio and the reconstituted particles rearrangement tested and characterized. Figure 1 shows that under our conditions, Wt rearranged mainly as three discrete particles of approximate Mw of 140, 440 and 670 kDa (Fig 1A). The relative quantification of the intensity associated with the bands confirmed a higher yield of the smallest population (Fig 1B). Instead, intra-chain crosslinked Wt (Wt+BS3) yielded a particle population, mainly represented by the largest complexes (indicated in Fig A by the black stealth arrow).

**Figure 1.**
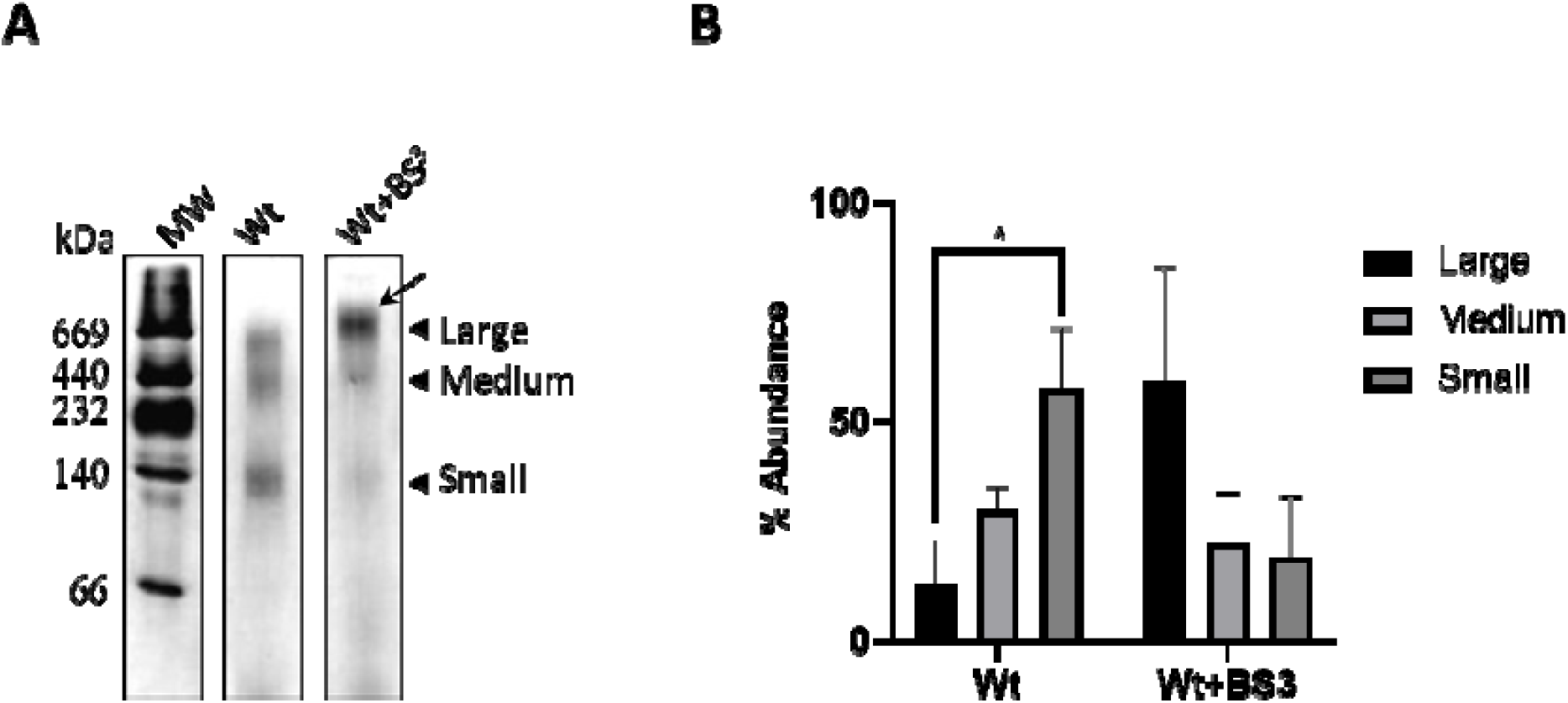
Characterization of sodium cholate-mediated reconstituted complexes. Wt and Wt+BS^3^ were incubated with DMPC MLV at a molar ratio 40:1 in the presence of sodium cholate as indicated in Methods. A) Home-made 4–30% non-denaturing, gradient gel electrophoresis (PAGGE) developed with silver staining. The apparent molecular weight was evaluated by comparison with high molecular weight commercial standards (labelled to the left). B) The relative amount of each population was estimated by quantifying intensity of the different bands in each lane. Bars represent media ± standard deviation of triplicates of three independent measurements as evaluated by t-test. Symbols * indicate significant differences at p ≤0.05.

In a different design, interactions of Wt with lipids may occur at the lipids transition temperature (Tm), where the apoA-I:phospholipid interaction was previously shown to be maximized[2]. The convenient Tm of the DMPC (24°C) is well suited to perform this test without requiring sophisticated lab heaters and avoiding proteins to be incubated under drastic conditions. In this regard, we set here to characterize the effect of intramolecular crosslinking by incubating Wt and Wt+BS^3^ with DMPC multilamellar vesicles (MLV) for 3 h at 24°C. First we analyzed the product of this interaction. As Figure 2 A shows, under these conditions large discrete-sized particles were obtained from Wt, with a low amount (around 20% as it is indicated in Fig 2B) of smaller complexes. Instead, and similarly to data shown for sodium cholate-mediated rHDL, Wt+BS^3^ yielded mostly one large population (indicated as in Fig 1A by the black stealth arrow).

**Figure 2.**
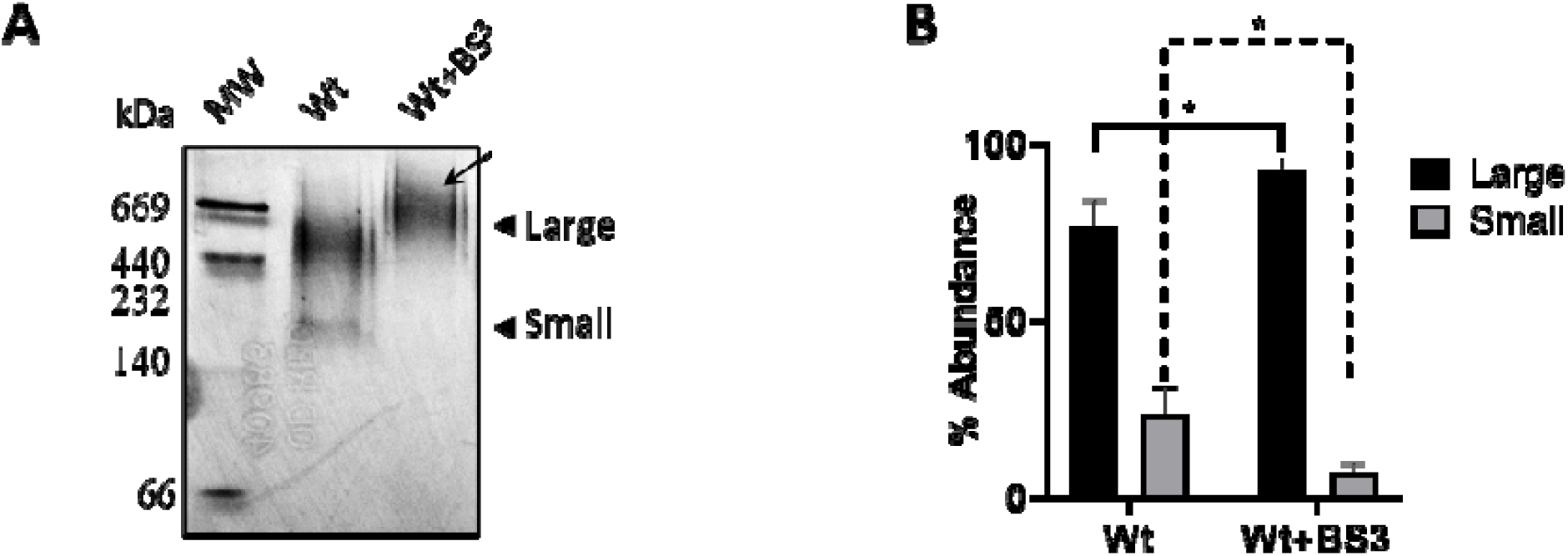
Characterization of the spontaneous interaction of Wt with DMPC. Multilamellar DMPC vesicles were incubated with Wt (0.05 mg/mL) or Wt+BS3 at a molar ratio 145:1 DMPC: protein at 24°C in phosphate saline buffer (PBS) pH 7.4 for 3 h (A) The relative size of the incubation product was estimated by 4–30% non-denaturing gradient gel electrophoresis (PAGGE) developed with silver staining as explained above. From left to right, commercial Mw marker, Wt and Wt+BS ^3^; B) The relative amount of each population was estimated by quantifying intensity of the different bands in each lane. Bars represent media ± standard deviation of triplicates of three independent measurements as evaluated by t-test. Symbols * indicated significant differences at p≤0.05 between Wt or Wt+BS3 large particles (continue line) or small particles (dashed line)

As the simplicity of this procedure makes it highly accessible to a vast biochemical research field, we further characterized its properties. First, we evaluated the importance of the regular buffers compositions used in a lab routine to perform this assay (either Tris or PBS) on the efficiency of the protein arrangement. As it is well known, the efficiency of the interaction may be followed by the decrease of the lipids turbidity as large MLV rearranges into small particles with lower light dispersion. From the registry of the absorbance during the incubation time, and as it was previously reported, Wt reorganized into small particles with fast kinetics (dark symbols in Fig 3 A). Instead, the lower efficiency of the crosslinked protein was evident as the kinetics of clearance was significantly diminished (white symbols). This behavior was independent of the buffer used during the incubation (either Tris A) or PBS B)) at a physiological pH. To better characterize the comparative clearance progression, Wt and Wt+BS3 absorbance was analyzed at different times. Even though the lower kinetics of the Wt+BS^3^ is observed since the beginning of the reactions, it became significant after 7.5-min incubation under these conditions (Fig 3C).

**Figure 3:**
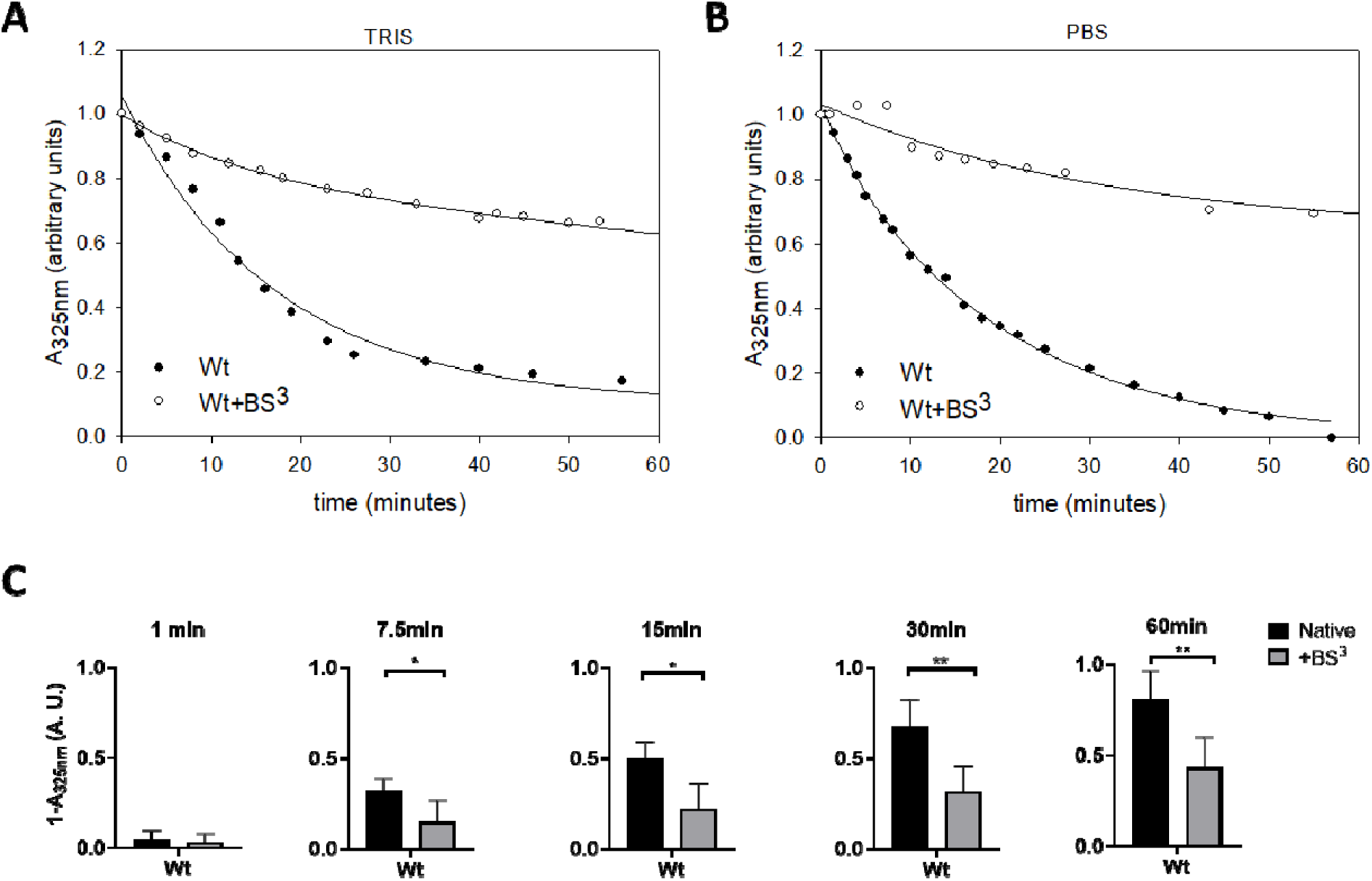
Analysis of the kinetics of DMPC clearance mediated by Wt. DMPC multilamellar vesicles were incubated with Wt (0.05 mg/mL) or crosslinked Wt (Wt-BS^3^) at a molar ratio 145:1 DMPC: protein at 24 °C, either in Tris 20 mM (A) or PBS (B) buffers, both at pH 7.4 for 1 h; Time dependence of the absorbance at 325nm was monitored during 1 h. C) Difference in the efficiency of micellization between Wt (black bars) or Wt-BS^3^ (grey bars) was estimated from the change in absorbance at 325 nm at each incubation time. Bars represent media ± standard deviation of triplicates of three independent measurements as evaluated by t-test. Symbols * and ** indicate significant differences at p ≤0.05 and p ≤ 0.01 respectively.

Next, we set to characterize the effect of temperature on the stability of the rearranged nanoparticles. When comparing the stability at 24°C, longer times (72 h versus 3 h) resulted in a small arrangement of Wt, yielding some degree of smaller particles (Fig 4A); instead, Wt+BS^3^ remained mainly as one discrete, large population (indicated by a black arrow).

**Figure 4.**
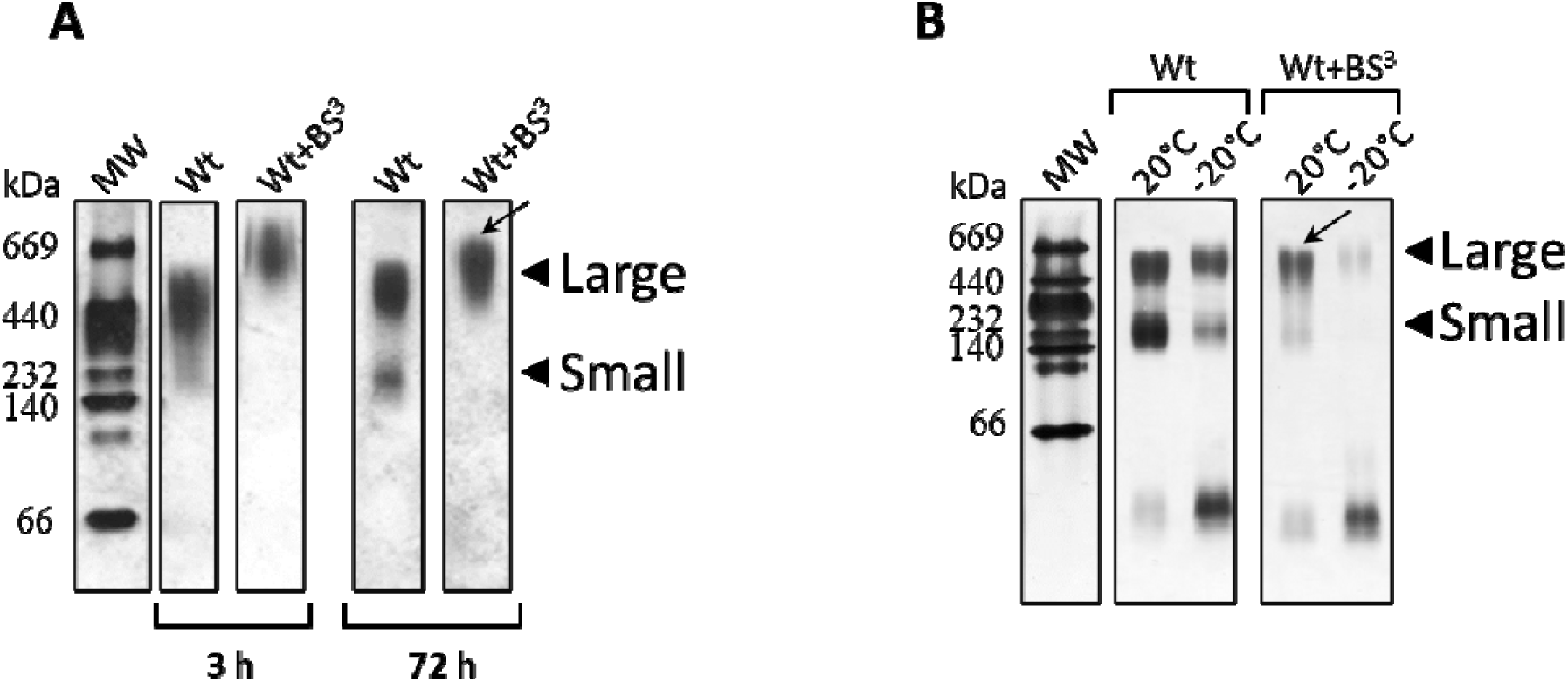
Stability of Wt complexes as a function of incubation time and temperature. DMPC multilamellar vesicles were incubated with Wt (0.05 mg/mL) or Wt+BS ^3^ at a molar ratio 145:1 DMPC: protein at A) 24 °C for 3 h or 72 h; B) Particles obtained after 72 h at 24°C are shown in A) were incubated for seven days either at 20 °C or −20 °C. The relative size of the incubation product was estimated by 4–30% non-denaturing gradient gel electrophoresis (PAGGE) developed with silver staining as explained above.

Afterward, we analyzed the stability of the rearranged particles by incubation at different temperatures. Particles obtained after 72 h at 24°C (Fig. 4A) were incubated for 7 days at 20°C resulting in a higher yield of the smaller particles and some amount of lipid-free protein for Wt. A lower effect was observed for Wt+BS^3^ on lipoprotein distribution (Fig 4B, labeled as above by the black stealth arrow). Taking lipid-protein complexes under freezing conditions strongly disrupted the weak interactions among Wt and lipids.

Finally, the different complexes obtained by incubation for 72 h at 24°C were isolated by size exclusion chromatography (Fig 5A), and reanalyzed afterwards by native PAGGE (Fig 5B). The most of the protein eluted as large complexes, especially in the case of Wt+BS^3^, which remained stable under the elution and concentrations steps, as shown in Fig 3B. The analysis of the complexes morphology by TEM indicated as expected that Wt formed the well-known disc-shaped complexes [1]. Crosslinked Wt also formed discoidal particles but with some elongated conformations.

**Figure 5:**
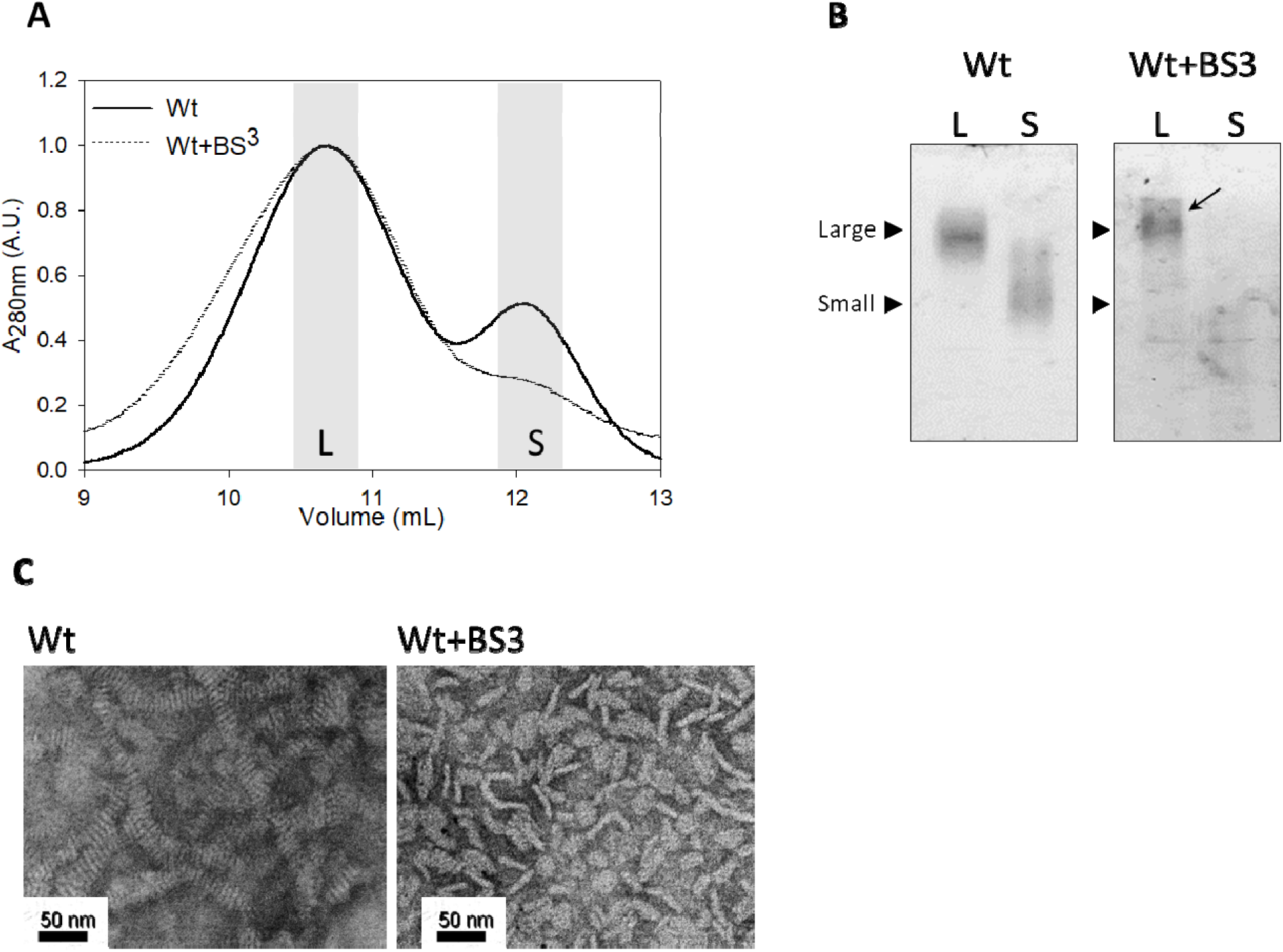
Characterization of apoA-I variants rearrangement. Wt and Wt+BS^3^ (at a protein concentration of 0.05 mg/mL) were incubated with DMPC for 72 h at 24.0°C at a molar ratio 145:1 and the product of the micellization analysed A) Size exclusion chromatography was performed through a Superose 6 HR column (Pharmacia), equilibrated with 50 mM Tris buffer pH 7.4 at a flow of 0.5 mL/min. Collected fractions corresponding to the main peaks (shaded in grey) were pooled, concentrated and analysed by a 4–30% non-denaturing, home-made gradient gel electrophoresis developed with silver staining B). Homogenous large sized Wt-BS^3^ complexes are labelled by a black stealth arrow; C) Morphology of the reconstituted HDL from Wt and Wt-BS^3^ was analysed by TEM under negative staining prior to FPLC isolation. Black bars in C) indicates the magnification scale used for the observations.

## Experimental Design, Materials and Methods

### Materials

Guanidinehydrochloride (GndHCl), cholesterol (Chol), sodium cholate, ethylenediaminetetraacetic acid (EDTA), sodium chloride (NaCl); dimyristoyl phosphatidylcholine (DMPC) was purchased from Avanti Polar Lipids. Alabaster, AL. His-purifying resin was from Novagen (Darmstadt, Germany). Bis- (sulfosuccinimidyl) suberate (BS^3^) and isopropyl-β-D-thiogalactoside (IPTG) were purchased from Thermo Scientific (Waltham, MA). All other reagents were of the highest analytical grade available.

### Protein variants purification

The cDNA of apoA-I with the native sequence was modified to introduce an acid labile Asp–Pro peptide bond between amino acid residues 2 and 3 of apoA-I, which allowed specific chemical cleavage of an N-terminal His-Tag fusion peptide. These constructs, inserted into a pET-30 plasmid (Novagen, Madison, WI), were transformed into BL21 (DE) Escherichia coli cells (Novagen, Madison, WI), then expressed by induction with IPTG and purified by elution through Ni-chelating columns (Novagen, Madison, WI) as described [3]. This resulted in a high yield of protein with a purity of at least 95% (determined by SDS-PAGE).

### Crosslinking of Proteins

Proteins were crosslinked at 0.05 mg/mL (monomeric state) in PBS pH 7.9 for 3 h without agitation at room temperature. Fresh BS^3^ was added (within 1 min after solubilization in PBS to avoid hydrolysis of free crosslinker) at 30:1 BS^3^:protein molar ratio[4]. Reactions were quenched by 15 min after the addition of Tris buffer to a 50 mM final concentration. The concentration of crosslinked proteins was calculated by the BCA Protein Assay Kit (Thermo Scientific (Waltham, MA)). The presence of the monomeric conformation after the treatments was confirmed by polyacrylamide gradient electrophoresis, either under native or denaturing conditions. Oligomers were not observed at these conditions. Proteins were dialyzed against PBS or TRIS buffer at pH 7.4, alternatively and stocked at - 20°C.

### DMPC multilamellar vesicles construction

Five mg of DMPC from a stock solution in chloroform was used to form a film in a round-bottom tube, dried by blowing a N_2_ atmosphere and exhaustively exposed to vacuum in a lyophilizer (Virtis) to evaporate the solvent. Then, PBS or Tris buffers pH 7.4 were added to a final DMPC concentration of 5 mg/mL. Multilamellar liposomes (MLV) were attained by extensive vortexing at room temperature for 5 min, heating at 37°C in three 30-sec cycles[5].

### DMPC Clearance assay

DMPC vesicles were added to the 0.05 mg/mL protein samples (pre-heated) until a final molar ratio of 145:1 DMPC:protein [2]. All reagents and instrument were pre-heated at 24°C. Samples were gently mixed (for 5 sec.) and clearance at 24°C was determined as light dispersion by monitoring absorbance (A) at 325nm in both spectrophotometers alternatively: a Helios Beta Single beam quartz coated (Thermo), and in a UV-visible Agilent Cary 8454. (Agilent Technologies). All DMPC experiments were performed in the presence of 0.05% sodium azide. Each datapoint was normalized to initial absorbance. Curves were adjusted by fitting to a double exponential decay in SigmaPlot software version 12.0. Efficiency of micellization (clearance) was determined as 1-A_325nm_ at fitted curves at indicated times. Five different experiments were used to determine averaged 1-A_325nm_ and standard deviation.

### Lipoprotein particles construction

#### By DMPC micellization

Same protein lipid ratio as mentioned above was used. Proteins at 0.05mg/mL were pre-incubated to a final temperature of 24°C (as the rest of the elements: cuvettes, lipids, tubes, etc). And lipids were added to proteins and gently mixed. Excess of lipids from tips or syringes needs to be removed prior to be added to the proteins. Incubations were performed always at 24°C varying the incubation time from 3 to 72 hours. Particle behaviour to temperature dependence was evaluated using particles obtained by 72 hours at 24°C. Stable particles were incubated at room temperature (20°C) or at −20°C, for 7 days. Samples were thawed at room temperature, and immediately seeded on gel for electrophoresis run.

#### By detergent rearrangement

Lipoprotein particles were constructed using the rearrangement method mediated by detergents [1]. Sodium cholate in PBS was added to DMPC MLVs to a final molar ratio of 40:60 (DMPC:sodium cholate). Once sodium cholate was added, initially “cloudy” DMPC MLV clarified by vortexing followed by 30-min incubation at 24°C. Lipids were added to proteins to final molar ratio 1:40:60 (protein:DMPC:sodium cholate) at 0.05 mg/mL, and samples were incubated overnight at 24°C. Detergent was removed by extensively dialysis of samples using at least 3 PBS batch changes for at least 3 h per bath, at 24°C (in a minimal relation of 1:1000 of sample to dialysis buffer).

### Polyacrylamide gel electrophoresis

Home-made native gradient 4-30% polyacrylamide gels were used to analyse lipoprotein particles size and relative amount. Each gel was imaged and transformed into an 8-bit image. If contrast or bright were modified, it has been applied to the entire gel. Densitometry and areas of curves in plots were obtained using ImageJ software version 1.51 j8. Data were normalized considering as 100% of density the sum of each lipoprotein band per lane. Triplicates were averaged and used to determine standard deviation for subsequent statistical analysis.

### Fast protein liquid chromatography (FPLC)

Lipoprotein particles were obtained by 72 h incubation of proteins at same DMPC:protein and protein concentration as described above. The relative size of the particles was estimated by size exclusion chromatography (SEC) using a Merck-Hitachi L6200 Intelligent pump. Samples were filtered in a 45 µm-pore syringe filter and injected (300uL each) through the Superose 6 HR 10/30 column, previously equilibrated with 50 mM Tris buffer pH 7.4 at a flow of 0.5 mL/min, and detected at 280 nm using a UV-VIS detector (Merck-Hitachi L4200). Points in curves were normalized to maximum of each dataset. Only collected samples being coincident to chromatograms peaks, were tested by native gradient electrophoresis.

### Transmission electron microscopy observations

Particle morphology was characterized by transmission electron microscopy (TEM) on a JEOL1200 EX. Samples were seeded on Formvar grids, contrasted with 0.5% phosphotungstic acid and visualized by negative staining. Lipoprotein samples were formed by DMPC incubation at 145:1 (DMPC:protein) at 0.05mg/mL protein concentration for 72 h at 24°C. Samples preparation for TEM imaging was performed at room temperature. Selected images are representative from 7 independent images captured from different grids zones. Bright and contrast were adjusted in ImageJ software version 1.51 j8 to improve the visualization of lipoprotein’s shape.

### Other analytical methods

For the statistical analysis, datasets were analysed in GraphPad Prism 8.0 software using parametric t-test with Welch’s correction or unpaired T-test corrected for multiple comparisons using the Holm-Sidak method. Only results with a significance level of p<0.05 were considered. Unless otherwise stated, the results either of biophysical assays were reproduced in three independent experiments and were reported as means of triplicates ± standard deviation.

## Ethics Statement

No animal or human samples have been used in this work

## Acknowledgments

This work was supported by the Consejo Nacional de Investigaciones Científicas y Técnicas (CONICET, PUE 22920160100002 to HG); Agencia Nacional de Promoción Científica y Tecnológica (ANPCyT, PICT-2016-0849 to MAT and PICT-2016-0915 to HG); Universidad Nacional de La Plata (UNLP) (M187 and M234 to MAT, PPID M014 to NAR).

## Authors’ individual contributions to the paper

Ivo Díaz Ludovico and Romina A. Gisonno: Conceptualization, Methodology, Writing; Validation; Horacio A. Garda, Marina C. Gonzalez, Validation, Conceptualization; Nahuel A. Ramella and M. Alejandra Tricerri, Conceptualization, Writing, Funding acquisition, Methodology.

## References

[1] C.E. Matz, A. Jonasf, Micellar Complexes of Human Apolipoprotein A-I with Phosphatidylcholines and Cholesterol Prepared from Cholate-Lipid, 257 (1982) 4535–4540.

[2] J.B.S. and B.C. Chang, Thermal Dependence of Apolipoprotein A-I-Phospholipid Recombination, Biochemistry. 19 (1980) 5637–5644.

[3] N.A. Ramella, O.J. Rimoldi, E.D. Prieto, G.R. Schinella, S.A. Sanchez, M.E. Jaureguiberry, M.S. Vela, S.T. Ferreira, M.A. Tricerri, Human apolipoprotein A-I-derived amyloid: Its association with atherosclerosis, PLoS One. 6 (2011). https://doi.org/10.1371/journal.pone.0022532.

[4] P. Cross-linkerst, N-Hydroxysulfosuccinimide Active Esters⍰: Bis (N-hydroxysulfosuccinimide) Esters of Two Dicarboxylic Acids Are, (1982) 3950–3955.

[5] H.J. Pownall, Q. Pao, D. Hickson, J.T. Sparrow, S.K. Kusserow, J.B. Masseyg, Kinetics and Mechanism of Association of Human Plasma Apolipoproteins with Dimyristoylphosphatidylcholine: Effect of Protein Structure and Lipid Clusters on Reaction Rates, (1981) 6630–6635.

